# The clinically relevant *MTARC1* p.Ala165Thr variant impacts neither the fold nor active site architecture of the human mARC1 protein

**DOI:** 10.1101/2022.03.26.485076

**Authors:** Michel A. Struwe, Bernd Clement, Axel J. Scheidig

## Abstract

Recent genome-wide association studies (GWAS) have shown a common variant of the human mARC1 protein to be associated with liver disease. Herein, we present the crystal structure of this mARC1 p.Ala165Thr at near-atomic resolution. No relevant differences between the structure of wild type mARC1 and the variant protein are detected.

## Results and Discussion

In the past years, advances in genome-wide association studies (GWAS) have provided novel insights into potential targets for treatment of diseases. In case of non-alcoholic fatty liver disease (NALF) and non-alcoholic steatohepatitis (NASH), various GWAS have identified the human mARC1 protein (encoded by the *MTARC1* gene) as a relevant contributor to these highly prevalent liver diseases.

Very recently, Schneider *at al*. have published a very impressive study on the effect of the p.Ala165Thr mutation of the *MTARC1* gene (Schneider *et al*., 2021). They found significantly reduced liver-related mortality in carriers of the p.Ala165Thr genotype when additional risk factors like type-2 diabetes were present. Remarkably, this effect was dose-dependent, meaning that a stronger protective effect was observed in homozygous carriers of the p.Ala165Thr allele compared to heterozygous carriers(Schneider *et al*., 2021).

A protective effect of the mARC1 p.Ala165Thr variant against liver disease and changes to lipid metabolism were first published in 2020 by Emdin *et al*. (Emdin *et al*., 2020, Emdin *et al*., 2021) and has since been confirmed by the work of other groups in additional studies (Luukkonen *et al*., 2020, Mann *et al*., 2020, Parisinos *et al*., 2020, Gao *et al*., 2021, Ghodsian *et al*., 2021, Janik *et al*., 2021). It has consequently been suggested, that mARC1 could present a target for future NALFD or NASH therapies (Bianco *et al*., 2021).

The strong association of the p.Ala165Thr variant with human disease was quite surprising to us, as various common variants of the mARC1 protein had been characterised *in vitro* some years ago, and no differences in either *N*-reductive activity towards a model substrate nor loading with the molybdenum cofactor were observed for recombinant proteins (Ott *et al*., 2014).

We have therefore now tried to determine, what effect the p.Ala165Thr amino acid exchange has on a protein-structure level. The mutation was introduced into our crystallisation construct by site-directed mutagenesis and protein for crystallisation was produced and crystallised essentially as described before (Kubitza *et al*., 2018). X-ray diffraction data were collected at the P14 beamline (Petra III, DESY, Hamburg), where crystals diffracted to near-atomic resolution (1.6 Å). The structure factors and model are available in the protein databank under the accession code 7P41.

When comparing the structure of the variant protein to the wildtype (PDB: 6FW2) (Kubitza *et al*., 2018), no relevant differences can be observed (**Figure 1A**). The molybdenum active site is well-defined (**Figure 1B**) by electron density and the overall fold of the protein is unaltered. The data suggest alternate conformations of Thr165 (Ala165 in the wildtype protein); however, successful mutation of this residue is obvious from the electron density (**Figure 1C**).

**Figure 1:**
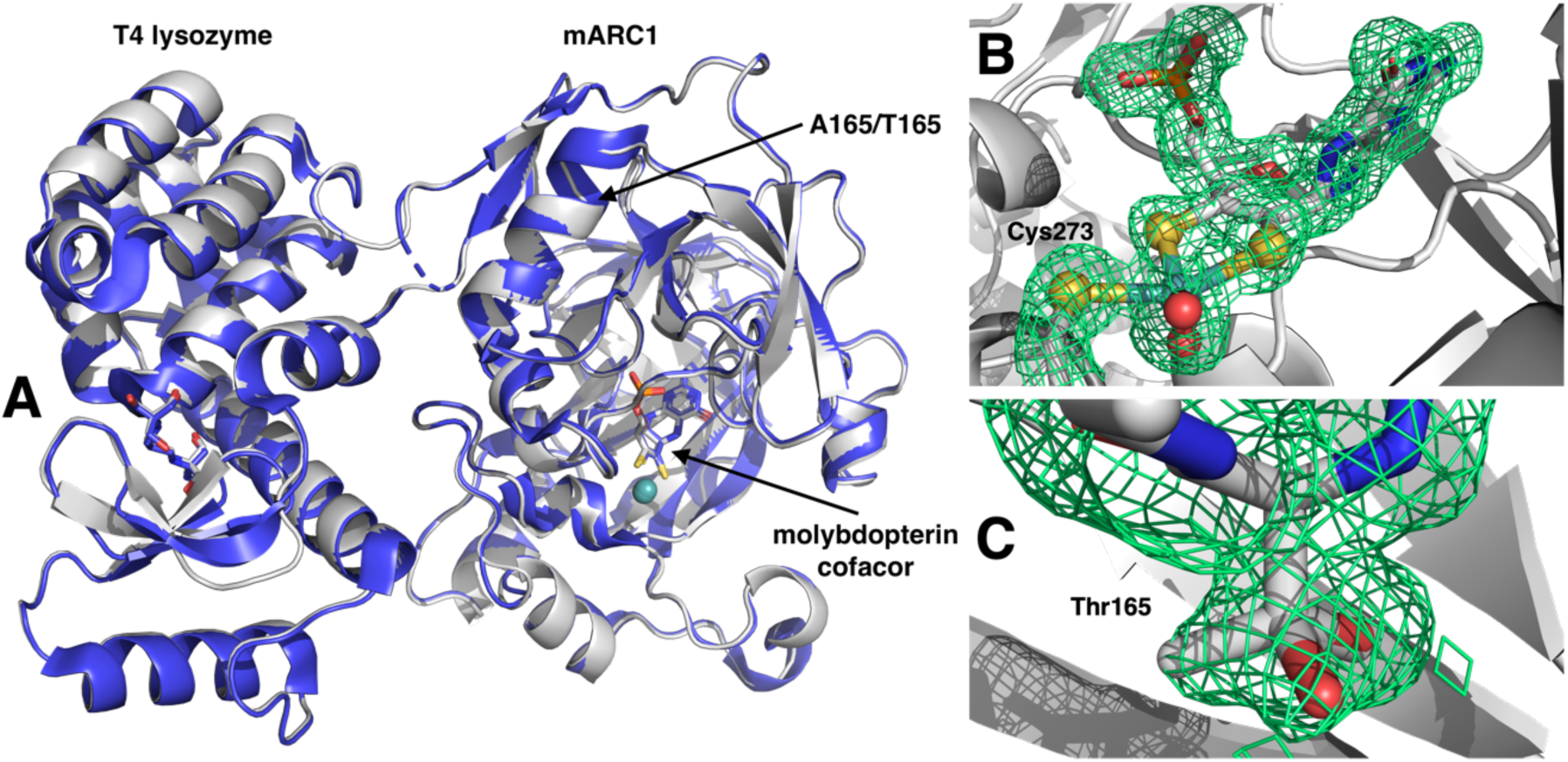
(**A**) Superimposed structures of both wildtype (blue) and variant (grey) mARC1 as fusion proteins with T4 lysozyme. The fold of both proteins is identical. (**B**) Electron density surrounding the molybdenum site clearly shows an intact pentacoordinate molybdenum site. (**C**) Thr165 with electron density indicating two conformations of Thr165. Electron density maps in panels B and C are *composite omit maps*, that were generated using simulated annealing to remove any model bias.

We have also conducted biophysical characterisations using differential scanning fluorimetry and could not detect relevant differences between the variant and wildtype proteins with respect to thermal stability over a wide pH-range (data not shown).

To summarise, structural and *in-vitro* characterisations of the p.Ala165Thr variant have failed to show any impact of the mutation on the protein with respect to activity, cofactor loading, structure and stability. While many have speculated, which potential endogenous substrate of mARC1 might be involved in liver disease, our data does not suggest any changes to the active site of the enzyme.

Thus, it must be questioned whether any known mARC1 activity is connected for the phenotype associated with the variant. Alternatively, a moonlighting activity of the protein, which is completely independent from the molybdenum active site and oxidoreductase activity could be responsible. Future work on the specific involvement of mARC enzymes in lipid metabolism and their interactions with other proteins in the cell will hopefully provide more insight.

## Supplementary information

### Supplementary information S1. Experimental details

#### Molecular biology

A plasmid for recombinant expression of the p.Ala165Tht-mARC1-T4 lysozyme fusion protein was generated from the plasmid previously used for expression of WT-mARC1-T4 lysozyme construct (Kubitza *et al*., 2018) using the QuikChange Lightning Site-Directed Mutagenesis Kit (Agilent #210518) according to the manufacturer’s instructions. Mutagenic primers were 5’-ccactgggcggtggcctcgccac-3’ and 5’-gtggcgaggccaccgcccagtgg-3’.

#### Protein production

Recombinant protein for crystallisation studies was prepared as described previously (Kubitza *et al*., 2018).

#### Crystallisation, data collection and processing

Crystals were prepared essentially as described previously (Kubitza *et al*., 2018) with minor modifications (see **Table S2**). X-ray diffraction data were collected on the P14 Beamline at Petra III, DESY, Hamburg (see **Table S3**). We used XDS (Kabsch, 2010) for indexing and integrating, *AIMLESS* for space-group determination, data scaling and merging (Potterton *et al*., 2018) and *PHASER* for molecular replacement (McCoy *et al*., 2007) with the previously published structure of mARC1 (PDB: 6FW2) as search model. The model was then refined by several cycles of automatic refinement within the *phenix* suite (Afonine *et al*., 2012) and manual corrections in *COOT* (Emsley *et al*., 2010).

#### Data visualisation

In order to compare the structures of WT mARC1 and the p.Ala165Thr variant, a structural overlay was generated using the *DALI* webserver (Holm, 2020). Molecular graphics shown in **Figure 1** were prepared in *PyMOL* (Schrödinger LLC).

**Supplementary Table S2.**
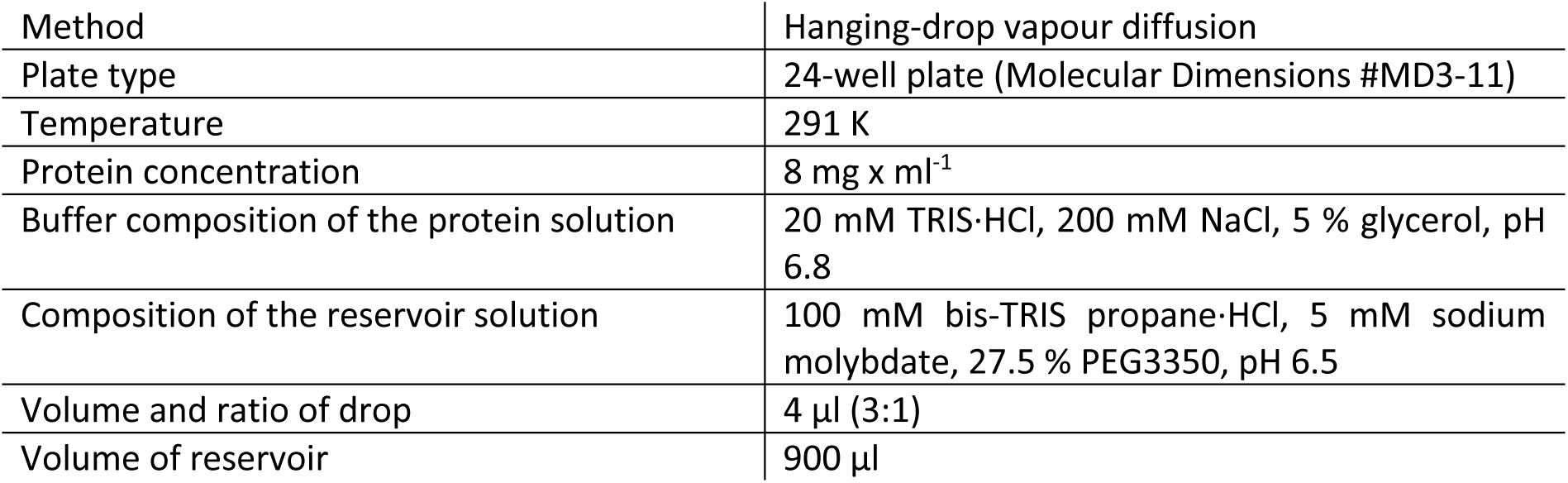
Crystallisation details

|  |  |
| --- | --- |
| Method | Hanging-drop vapour diffusion |
| Plate type | 24-well plate (Molecular Dimensions #MD3-11) |
| Temperature | 291 K |
| Protein concentration | 8 mg x ml<sup>-1</sup> |
| Buffer composition of the protein solution | 20 mM TRIS-HCl, 200 mM NaCl, 5 % glycerol, pH 6.8 |
| Composition of the reservoir solution | 100 mM bis-TRIS propane-HCl, 5 mM sodium molybdate, 27.5 % PEG3350, pH 6.5 |
| Volume and ratio of drop | 4 µl (3:1) |
| Volume of reservoir | 900 µl |

**Supplementary Table S3.** Data collection.

|  |  |
| --- | --- |
| Diffraction source | PETRA III, EMBL c/o DESY beamline P14 (MX2) |
| Wavelength | 0.9762 Å |
| Temperature | 100 K |
| Detector | Dectris pilatus 6M pixel |
| Rotation per image | 0.1 ° |
| Total rotation range | 360 ° |
| Exposure time per image (s) | 0.01 s |
| Space group | $P2_12_12_1$ |
| a, b, c | 61.06 Å, 74.89 Å, 111.16 Å |
| $\alpha$, $\beta$, $\gamma$ | 90 °, 90 °, 90 ° |
| Mosaicity | 0.13 |
| Resolution range | 53.52 – 1.60 Å (1.63 – 1.60 Å)* |
| Total N of reflections | 910,173 (44,210)* |
| N of unique reflections | 66,911 (3,234)* |
| Completeness (%) | 98.6 % (97.3 %)* |
| Redundancy | 13.6 (13.7)* |
| Average $I/\sigma(I)$ | 21.0 (9.1)* |
| $R_{r.i.m.}$ | 0.096 (0.248)* |
| $R_{p.i.m.}$ | 0.026 |

**Supplementary Table S4.**
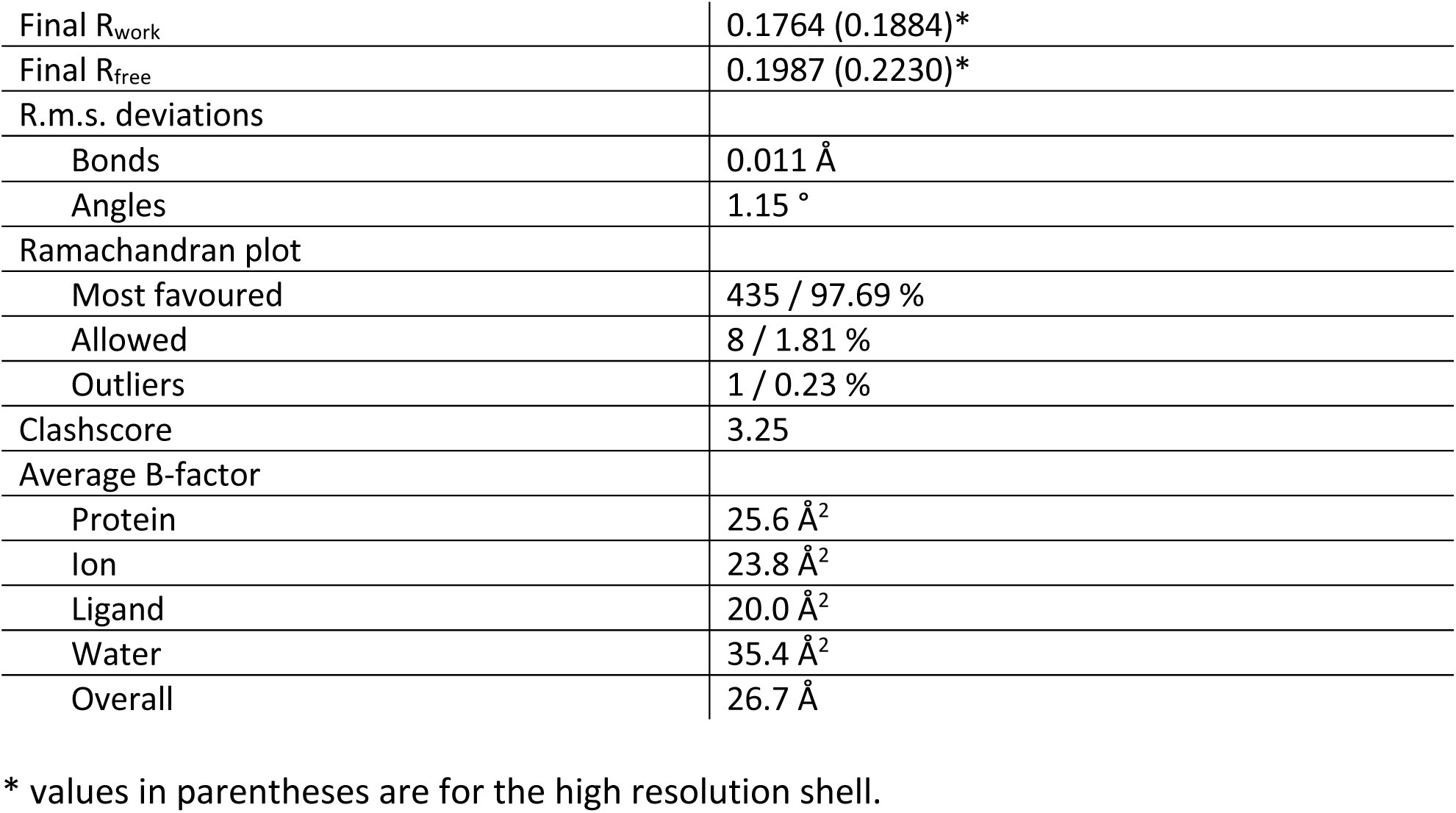
Refinement statistics.

